# Cannabinoid type-1 (CB_1_) receptors in glial cells promote neuromuscular junction repair following nerve injury

**DOI:** 10.1101/2024.01.12.575382

**Authors:** Roberta Piovesana, Sophie Charron, Danielle Arbour, Giovanni Marsicano, Luigi Bellocchio, Richard Robitaille

## Abstract

Cannabinoids are frequently used in the treatment of neuropathic pain related to nerve injury. However, despite evidence for their roles in the regulation of axonal guidance and synapse formation during development of the central nervous system (CNS), their possible involvement in response to peripheral nerve injury remains poorly defined and the knowledge of its role is mostly related to the peripheral sensory system.

Following nerve injury, contemporary to axonal repair, massive morphological and functional changes reshape synaptic elements at neuromuscular junctions (NMJs) aiming to promote their reinnervation. This process is mediated in part by Perisynaptic Schwann cells (PSCs), glial cells at the NMJ essential for its maintenance and repair.

Here we investigated the novel role of Cannabinoid type-1 receptor (CB_1_R) at NMJ, in particular on PSCs, during motor nerve recovery following nerve injury. Using morphological analysis, we studied the consequences of CB_1_R pharmacological and genetic blockade following denervation and reinnervation in adult NMJs. CB_1_R blockade caused an acceleration of the denervation process followed by a great delay in reinnervation as indicated by a significant percentage of denervated NMJs, accompanied by a decrease of mono- and poly-innervated NMJs. Remarkably, a similar phenomenon was observed when CB_1_R is selectively knocked-out in glia, indicating that the protective actions of these receptors are largely glia-dependent.

These data highlight a novel role of the endocannabinoid system at NMJs, where the CB_1_Rs on PSCs can control NMJ denervation and reinnervation following nerve injury. A better understanding of the functional mechanisms underlying CB_1_R role in NMJ repair may contribute to finding a new pharmacological treatment having a dual role in improvements of motor recovery and in pain-related relief.

## 1. Introduction

Peripheral nerve injuries are a common clinical problem, usually consequences of trauma or secondary to other conditions, such as tumour surgical excision. Although the peripheral nervous system (PNS) can regenerate, its ability is impacted by the age of the patient, type of injury and its proximity to the nerve cell body^1^. Moreover, the time of regeneration is crucial; delays in repair and/or intervention impede adequate sensory and motor recovery^2^.

No advances in the clinical treatment for nerve injury have been made in the last 30 years and its impact on the patient’s life (e.g. motor recovery, related pain) is often underestimated. Also, efficient pharmacological treatments are not available and often solely targeting pain-related relief^1,3^. One of the possible targets is the Endocannabinoid system (EC), known to be present on the sensory system and used in the treatment of neuropathic pain^4–6^, also associated with nerve injury^7–9^.

The EC, through its CB_1_ (CB_1_R) and CB_2_ (CB_2_R) receptors, modulates many functions involved in numerous physiological and pathological conditions^10–14^ in PNS and central nervous system (CNS) ^15–23^. The main characteristic of CB_1_R is its broad distribution in many different cell types, including neuronal and glial cells, and also in different intracellular compartments such as mitochondria^12^. In CNS, CB_1_R is expressed from the earliest stages of the embryo development and follows neuronal differentiation where it is involved in axonal growth. CB_1_R pharmacological blockade in the early embryo developmental stage is detrimental for axon pathfinding and fasciculation^24^. While CB_1_R inhibition of cultured hippocampal neurons at very early developmental stages had no effect on axonal growth, cannabinoid administration promoted this process. By contrast, subsequent dendritic growth is compromised by CB_1_R inhibition^25^. Interestingly, endocannabinoids are also able to regulate glial cell development and maturation. For instance, microglia express cannabinoid receptors, produce endocannabinoids and express enzymes for their degradation^26,27^; furthermore, enhancing endocannabinoids signaling has been shown to exert an anti-inflammatory effect and induce neuroprotective microglia phenotype^27^. Astrocytes express functional CB_1_Rs; cannabinoids local application evoked astroglial Ca^2+^ increase in the hippocampal tripartite synapse via CB_1_ receptor activation as confirmed by the blockade by the CB_1_R selective antagonist AM251^28,29^. Moreover, enzymes responsible for the synthesis of the endocannabinoid were found in oligodendrocytes at different developmental stages and the CB_1_R and CB_2_R blockade impaired their differentiation into mature oligodendrocytes^30–32^.

CB_1_R expression has been found at Neuromuscular Junctions (NMJs) of lower vertebrates where it co-localizes with ACh receptors^33^, but there is yet no evidence of CB_1_R expression and function in Perisynaptic Schwann cells (PSCs) level, glial cells essential for NMJ maturation, synaptic efficacy and plasticity and repair^34–40^. Indeed, nerve injury induces changes not only at the axonal level but also at NMJs where morphological and physiological changes are observed during the processes of denervation and reinnervation^41–44^. These events are strictly regulated by the PSCs ability to detect synaptic activity through acetylcholine muscarinic receptors (mAChRs) and purinergic receptors in a Ca^2+^ dependent manner^35–37,41,45–49^. PSCs are able to switch into a repair phenotype when mAChRs activation is reduced^41,42,48,50,51^. Importantly this happens following sciatic nerve crush, during synapse formation and aging^42,48,51^. Altogether these observations indicate that PSCs are essential in the maturation, maintenance and in the repair of NMJs following nerve injury^41^. However, while changes in pre- and post-synaptic elements have been deeply investigated, comprehensive knowledge underlying PSCs repair regulation is still poorly defined.

Despite an impressive amount of knowledge of the diverse effects of CB_1_Rs on axonal growth in CNS, its roles in the PNS and at the NMJ are ill defined, limiting our knowledge on the sensory system and its potential therapeutic use is exclusively in the treatment of neuropathic pain^4–6^, also associated to nerve injury^7–9^. Here, we investigated if the PSCs’ ability to regulate nerve repair response could be mediated by CB_1_R and, thus, its selective activation influences NMJs reinnervation. We tested CB_1_R role in motor recovery following nerve injury. To address this uncharted question, we used mouse nerve-muscle preparations to evaluate whether CB_1_R blockade or activation can regulate NMJ denervation and reinnervation following motor nerve injury. We found that CB_1_R blockade facilitated NMJ denervation induced by nerve injury and that it also significantly delayed the repair process in a glial-dependent manner. Importantly, *in vivo* CB_1_R activation, through the agonist WIN 55, 212-22, potentiated motor recovery already after 16 days post-injury, increasing the presence of mono-innervated NMJs.

Hence, we present a new CB_1_R-mediated mechanism in glial cells at NMJs, which enhances NMJ proper reinnervation following injury.

## 2. Methods

### 2.1 Mouse models

Complete nerve crushes were performed in adult male mice homozygous for the motor neurons expression of the jellyfish yellow fluorescent protein (YFP) under the control of the Thy1 promoter (B6.Cg-Tg(Thy1-YFP)16Jrs/J stock number 003709; The Jackson laboratories; referred as Thy1 mice) at 120-180 postnatal days as previously described^41^. Two CB_1_R knock-out mouse models were also used: the full CB1-KO^52,53^ and a conditional mouse mutant model in which CB_1_R is selectively knocked-out from cells expressing the glial fibrillary acidic protein (GFAP), such as astrocytes and PSCs. GFAP-CB1-KO mice were generated using the CRE/loxP system^54^. Briefly, mice carrying the ‘‘floxed’’ CB1 gene (CB1f/f)^55^ were crossed with GFAP-CreERT2 mice^56^, using a three-step backcrossing procedure to obtain CB1f/f;GFAP-CreERT2 and CB1f/f littermates, called GFAP-CB1-KO and GFAP-CB1-WT, respectively. CreERT2 protein is inactive in the absence of tamoxifen treatment^56^. Deletion of the CB1 gene was obtained in adult mice (7-9 weeks-old) by daily intraperitoneal (IP) injections of tamoxifen (1 mg, 10 mg/mL in 90% sesame oil, 10% ethanol) for 8 days. Mice were used 3-5 weeks after the last tamoxifen injection^54^. This strategy avoids consecutives problems of neuronal contamination using GFAP promoter during development^56^. All experiments were performed in accordance with the guidelines of the Canadian Council of Animal Care and the Comité de déontologie de l’expérimentation sur les animaux of Université de Montréal, the Committee on Animal Health and Care of INSERM, the French Ministry of Agriculture and Forestry (authorization number 3306369) and the French Ministry of Higher Education Research and Innovation (authorization APAFIS#33548). Only males were used in the present study. Mice had *ad libitum* access to fresh water and food.

### 2.2 Nerve Injury

Mice were anesthetized using 2% isoflurane in 97%–98% O_2_ delivered through a mask. The skin in the posterior lower back of the trunk was shaved and washed with a mix of 70% alcohol and povidone iodine solution. A small skin incision was made and the biceps femoris and gluteus maximus were gently separated from each other to expose the sciatic nerve. The sciatic nerve was then crushed once for 15 s using a Moria micro-serrated curved forceps (MC31) to ensure a complete denervation. The skin was then closed with sutures and animals were kept warm to prevent hypothermia. Contralateral leg underwent the same procedure except for the nerve crush manoeuvre (Sham). Buprenorphine, (0.1 μg g^-1^), was applied before the surgery, 6–8 h after the surgery, and the following day. This procedure minimises muscle inflammation since the injury was made close to the hip, far away from the NMJs, and gives the operator the time to study different phases of the process. Denervation process is complete within 24 h after nerve injury, and reinnervation takes place 12 to 18 days post injury. Hence, EDL (*Extensor digitorum longus*) muscles were collected after 18-48 hours post-injury (hpi) for denervation studies to evaluate the ongoing denervation (18 hpi) and the complete denervation (48 hpi) and muscles were collected 16 days post-injury (DPI) for reinnervation evaluation.

### 2.3 Drug treatments

CB_1_R agonist (WIN 55,212-2; 3 mg/kg or 0.3 mg/kg) and antagonist (AM251; 3 mg/kg) were delivered by intraperitoneal (IP) injections. For the denervation process, injections were performed two days before and on the day of the surgery. For the reinnervation process, injections were performed daily during the synaptic competition events^41^, between 12- and 16 DPI. Drugs were diluted in 7.7% DMSO, 4.6% Tween 80 in saline (vehicle). Control mice were injected with the vehicle solution. Treatments were performed at the same time of the day (1 PM ± 1 h) and mice activity was monitored for 2 h after treatments.

NMJs of controlled, contralateral muscles, treated with vehicle, AM251 or WIN 55,212-2, were unaltered, confirming that vehicle or antagonist did not have any impact on healthy, not-nerve injured NMJs (*supplemental data*).

### 2.4 Protein extraction and Western blot

EDL muscles without tendons and hippocampi as positive controls were rapidly collected and frozen in liquid N_2_ and stored at -80 C until protein extraction. Samples were then weighted to calculate the volume of lysis buffer to be used (4.5 x weight; 50 mM Tris-HCl pH 7.5, 1 mM EDTA pH 8, 150mM NaCl, 1% triton-X and 1:10 protease inhibitor (Sigma Fast inhibitor tablet; S8820; Sigma-Aldrich, Saint Louis, USA). Samples were maintained on ice and partial mechanical digestion was done. Samples were then vortexed for 15 s and incubated for 30 min on ice before a second mechanical digestion was performed. SDS/NP40 (0.5X of the weight of the original tissue) were added and samples were vortexed for 15 s and incubated for 10 mins at Room Temperature (RT). Afterwards, samples were centrifuged for 30 min at 14.8K rpm at 4°C. Surnatant (protein extract) was then collected, and protein concentration was determined using Pierce^TM^ BCA Protein Assay Kit (Thermo Fisher Scientific, Waltham, MA, USA), according to the manufacturer’s protocol.

Sample buffer (4×, 240 mM Tris-HCl pH 6.8, 8% (*w/v*) SDS, 40% (*v/v*) Glycerol, 0.1% (*w/v*) Bromophenol Blue, 10% (*v/v*) β-mercapoethanol, pH 8.3-8.8) was added to 30 μg of protein lysates. Heat-step was not performed for CB_1_R detection. Samples were loaded onto 4-15% Mini protean tgx-stain-Free gels (Bio-Rad Laboratories Canada Ltd, Ontario, CA, Cat#4568083) and run at 100-120V using Tris-glycine running buffer [25mM Tris, 190mM glycine, 1% (w/v) SDS]. Resolved proteins were transferred for 60 min onto PVDF blotting membranes (Thermo Fisher, Ca#: 88518) at 100V at 4 C, in transfer buffer (250 mM Tris-base; 192nM glycine, 10% (*v/v*) methanol; pH 8.3). In order to confirm successful protein transfer, membranes were stained with Ponceau red (Sigma-Aldrich, Poole, UK), before being blocked for 1 h in a Tris-buffer saline (TBS)-Tween Solution (10mM Tris pH 7.5, 100mM NaCl, 0.1% (*v/v*) Tween) containing 5% (*w/v*) of non-fat dry milk (blocking buffer). Membranes were incubated with the primary antibody (CB_1_R, 1:500 Abcam #23703; GAPDH, 1:5000, Cell signaling #21185) diluted in blocking buffer, overnight at 4 C. Membranes were then washed 3 times for 10 mins with TBS-Tween buffer and incubated for 1 h at RT with anti-rabbit (HRP) secondary antibody (1:20000, Ca# 711-035-152; Jackson ImmunoResearch Lab, Pennsylvania, USA) for chemiluminescence detection. To determine housekeeping protein (GAPDH), membranes were stripped with a glycine solution (100mM, pH 2.9; Sigma-Aldrich, UK) for 15 min at RT and, after three washes, blocked again for 1 h in 5% (*w/v*) of non-fat dry milk prior to further blotting.

Membranes were exposed to SuperSignal West Pico Chemiluminescent Substrate (Thermo Fisher Scientific, Waltham, MA, USA; Ca# 34580) for signal detection. The optical density (OD) of each protein band was analysed with ImageJ imaging software (Version 1.52a, National Institutes of Health NIH, United States) and normalized against the OD of the protein reference band.

### 2.5 Immunohistochemical labeling of NMJs

Immunohistochemical labeling of NMJs was performed on whole mount preparations to visualize the three components of the NMJ (PSCs, presynaptic terminal and postsynaptic endplate) as previously described^46,48,57^. Briefly, muscles were pinned in Sylgard-coated dishes and fixed with 4% formaldehyde (PFA; Mecalab, Canada) at RT for 10 min, rinsed three times with Phosphate Buffered Saline (PBS) and then permeabilized with 100% cold methanol for 6 min at -20 C. For the blockade of non-specific labeling, muscles were incubating in a solution containing 10% normal donkey serum (NDS; Jackson ImmunoResearch Laboratories) and 1% Triton X-100 for 30 min at RT. PSCs were then labeled with a S100 rabbit antibody (1:4, DAKO) for 2 h at RT. After rinsing three times with PBS containing 0.01% Triton X-100, presynaptic terminals were labeled using two primary antibodies, a chicken anti-neurofilament M (1:2000, 212-901-D84, Rockland) and a mouse IgG1 anti-synaptic vesicular 2 (1:2000, AB_2315408, DSHB) for 2 h at RT. Secondary antibodies (Alexa488 α-chicken/Alexa 488 α-mouse IgG1, Alexa 647 α-rabbit IgG, 1:500, Jackson ImmunoResearch) were incubated together for 1 h at RT. Muscles were then rinsed and incubated with α-BTX (Alexa594, 1:500, ThermoFisher) for 45 min at RT. All antibodies and α-BTX were diluted in PBS containing 1% Triton X-100 and 2% NDS. Whole muscles were mounted in a Prolong Gold antifade medium or Prolong Diamond medium to preserve endogenous fluorescence of the YFP protein (Molecular probes by Life technologies). All three elements of the NMJs were simultaneously visualized using FV1000 Olympus confocal laser-scanning microscope.

### 2.6 Morphological analysis

NMJs are classified in three groups according to the state of their innervation. For each NMJ, the presence of presynaptic elements (NFM-SV2) over the nAChRs α-BTX-labeled endplate was determined. During denervation experiments (18 hpi), NMJs were classified in 3 categories (supplementary data, Table 1): *early ongoing denervation* (called ‘early ongoing’; less than 30% of denervated NMJ area), *denervating NMJ* (up to 70% of denervated NMJ area) and *denervated NMJ* (70% or more of denervated NJM area). For reinnervation process (supplementary material, table 2), NMJs were classified in four categories: 1) *denervated NMJs* (no nerve terminal label on NMJs), 2) *mono-innervated NMJs* (partial or full coverage by a single axon) or 3) *poly-innervated NMJs* (at least two independent axons). It is important to note that complete innervation at 16 DPI is very rare (about 2% of NMJs^41^). NMJs were counted as mono- or poly-innervated when the coverage of a single axon or multiple axons were over 60% of the total end-plate area (Btx staining). For a better understanding and examples, see table 2 in supplementary material.

### 2.7 Statistical analyses

All statistical tests were performed in GraphPad Prism software (version 9.3.0) and results are presented as mean ± SEM. Mann–Whitney test was used to compare two different conditions (Vehicle vs treated muscles, WT/ GFAP-CB1-WT vs full-CB1-KO/GFAP-CB1-KO). The confidence level was set at 95% (α=0.05). Analyses were deemed significant at p <0.05. For the sample size, “N” represents the number of muscles from different animals (biological replicate number) and “n” the number of NMJs (number of observations).

## 3. Results

CB_1_Rs regulate axonal guidance and growth and synapse formation during CNS development but there is little information regarding their relevance for NMJ functions and their possible involvement on NMJ motor recovery following injury remains unexplored. Owing to the central role of PSCs in the regulation of NMJ maintenance, function, and repair, we hypothesized that CB_1_R could regulate NMJ repair in a PSCs dependent manner. Thus, we investigated the functional NMJ recovery using the pharmacological inhibition and global and glial genetic ablation of CB_1_R. Moreover, with a therapeutic perspective of the endocannabinoid, we explored the activation of CB_1_Rs using agonist WIN 55,212-2.

### 3.1 CB_1_R expression at NMJs and in PSCs

While CB_1_R expression was reported along the muscle fibers and at the endplate, co-localized with ACh receptors^33^, there are no data showing CB_1_R expression in PSCs. Thus, we firstly performed a western blot analysis showing that CB_1_R were present in muscles (Fig. 1A). No protein bands were found in the CB_1_R-KO mice. Hippocampal protein extract was used as positive control.

**Fig. 1.**
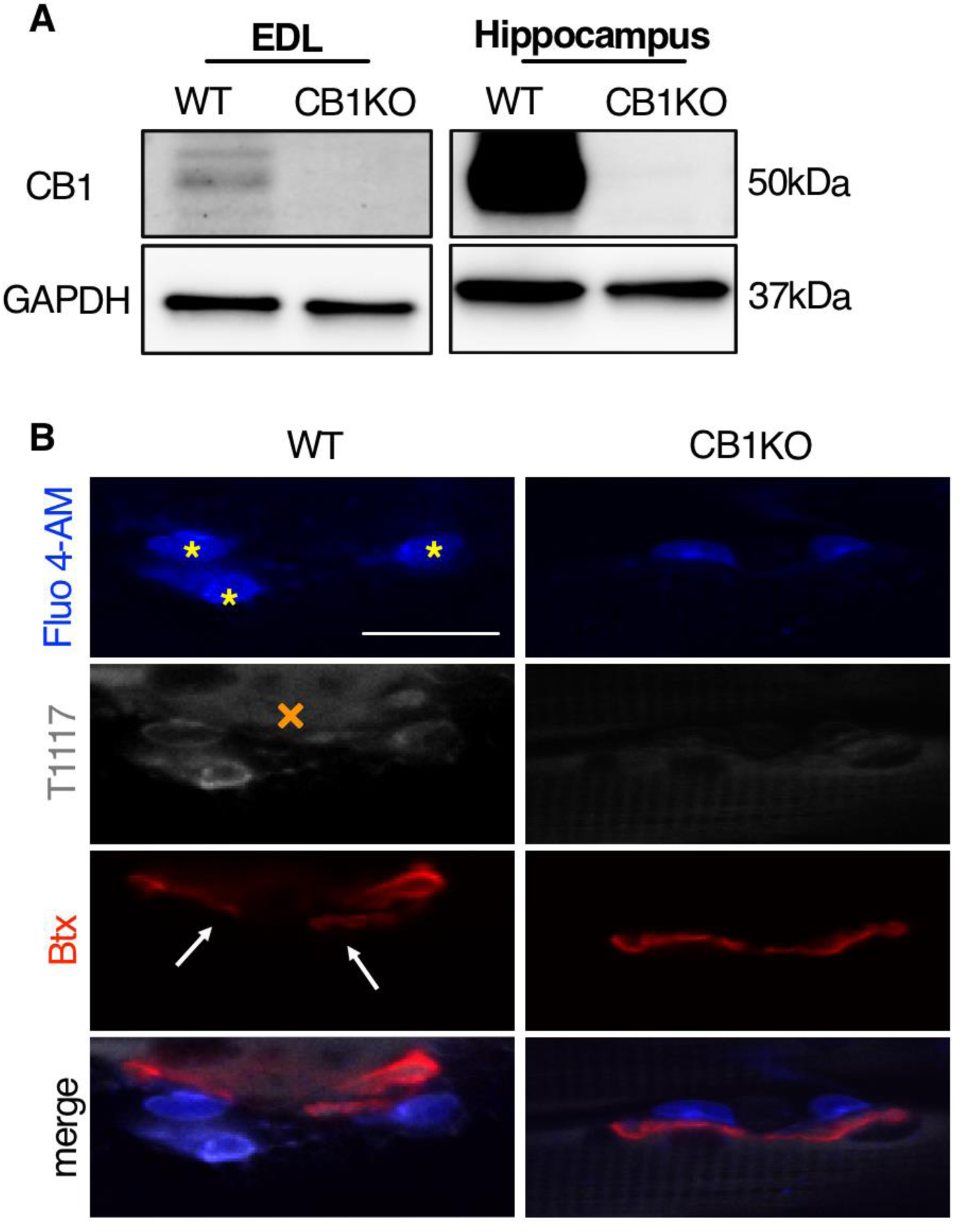
Expression of CB_1_R at NMJ. (A) Western blot of CB_1_R protein in EDL muscles from WT and full-CB_1_-KO mice. Hippocampus has been used as positive control. (B) PSCs loaded with fluo4 AM (blue), labeled with T1117, the fluorescent analogue of AM251 (grey) and nAChRs labeled with Btx (red, white arrow). Note that T1117 labelled muscle fibers (orange X) and PSCs (yellow asterisks) in WT mice. No T1117 labeling was observed in full-CB_1_-KO mice. Scale bar = 20μm.

The availability of good CB_1_R antibodies for immunostaining is very limited and they often show significant unspecific staining in full CB_1_-KO owing to the high homology of CB_1_R with other GPCRs (e.g. CB2R, GPR55). Hence, to circumvent this problem, we used T1117, a fluorescent analogue of the CB1R antagonist AM251. T1117 (100 nM) was bath applied to *ex vivo* nerve-muscle preparations for 30 min. As shown in Figure 1B, PSCs were labeled alongside the post-synaptic area which is consistent with the CB_1_R expression in muscle. Importantly, no labeling was observed in CB1-KO mice (Fig. 1B).

### 3.2 CB_1_R blockade accelerates denervation following nerve injury

We first tested whether CB_1_R regulates NMJ denervation following injury. Within 24 h following nerve injury, NMJs undergo a denervation process leading to the clearance of the nerve terminal from the injured axon, clearing the basal lamina in preparation for the reinnervation process controlled by axonal Schwann cells and PSCs^42,58,59^. A failure of the denervation process compromises the following NMJ reinnervation and subsequent atrophy of the muscles with a marked decrease in contractile force production^60^. Hence, we hypothesize that CB_1_R blockade will be deleterious on the denervation process.

During the denervation process, NMJs were found in different stages of denervation (Table 1 as reference). After 18 hpi, we observed an increased percentage of NMJs at advanced stages of denervation (denervated NMJs) when CB_1_R were blocked by *in vivo* treatment with AM251 (Fig. 2A, B; (* p < 0.05; N = 6; n = 91). Increased CB_1_-mediated denervation was confirmed using the full CB_1_-KO mice (Fig. 2A and C; ** p < 0.01; N = 5 n = 69). Moreover, using the GFAP-CB_1_-KO to selectively knock-out CB_1_R from glial cells, we also observed a significant increase of the percentage of denervated NMJs within 18 hpi (Fig. 2D; * p < 0.05; N = 5, n = 72), consistent with the results observed in general CB_1_-KO. These results suggest that the lack of CB_1_R on PSCs caused the acceleration of the denervation process.

**Fig. 2.**
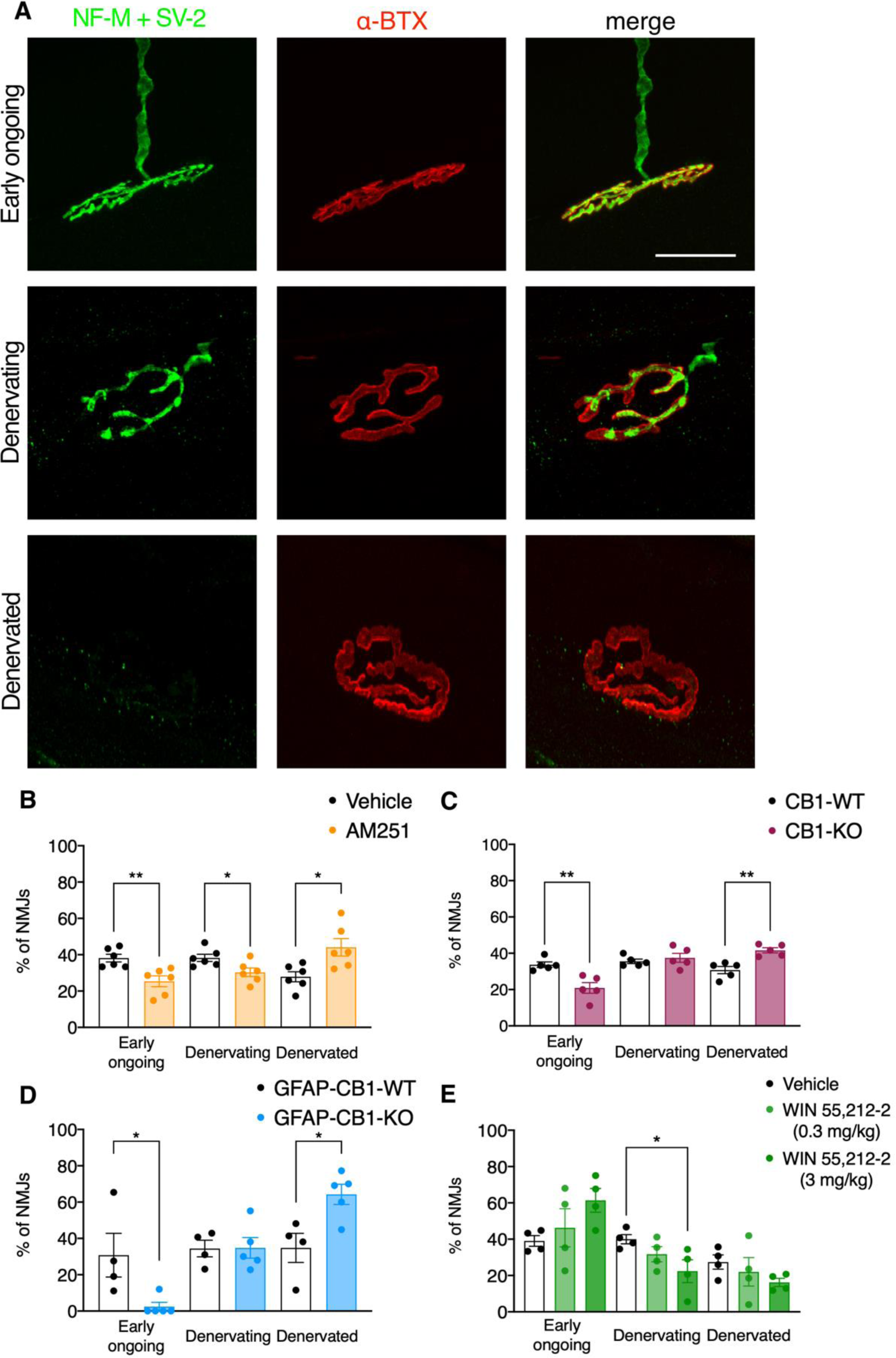
CB_1_R antagonist accelerates the denervation following nerve injury. A) Representative images of innervation states 18 h post-injury; (presynaptic, SV-2/NFM, green; postsynaptic, α-BTX, red). B) CB_1_R blockade through the treatment with AM251 (3 mg/kg) induced a significant decrease in early ongoing denervated NMJs (** p < 0.01; N = 6; n = 55) and a significant increase of denervated NMJs (* p < 0.05; N = 6; n = 91). C) Denervation was also accelerated in full CB_1_-KO (** p < 0.01; N = 5 n = 69) and GFAP-CB_1_-KO (D; (* p < 0.05; N =5, n = 72). E) WIN 55,212-2 treatment (0.3 or 3 mg/kg) did not alter the denervation process (Fig. E; ns; N = 4). Scale bar = 30 *μ*m. Results are presented as mean ± SEM.

To test if the activation of CB_1_R could alter the normal denervation process following nerve injury we tested the agonist WIN 55,212-2. Since CB_1_R agonists are notorious to often exert biphasic effects (i.e. having opposite effects depending on the concentration)^61–63^, two doses of WIN 55,212-2 were tested (3 mg/kg or 0.3 mg/kg). NMJ morphological analysis at 18 hpi revealed that, at these doses, activation of CB_1_R did not have any effect on the denervation process (Fig. 2E; ns; N = 4). Overall, these data show that blockade of CB_1_R accelerates the NMJ denervation while its activation did not have any effect.

### 3.3 CB_1_R absence is detrimental for reinnervation

The ultimate goal of nerve injury response and repair is the reinnervation of the NMJs and a functional motor recovery. To determine if CB_1_R contributes to NMJ innervation following injury, we studied NMJ reinnervation 16 DPI in mice treated daily with the CB_1_R antagonist AM251 during the synaptic competition period as recently determined (12-16 DPI)^41^. We previously showed that, similar to their postnatal maturation^48,51^, NMJs undergo a complex synaptic competition and show a similar percentage (∼ 50%) of mono- and poly-innervated at 16 DPI^41^. Similar results were independently obtained in control groups on different mice (Thy1 vehicle-injected) and WT littermates of CB_1_-KO and GFAP-CB_1_; Fig. 3B, C, D), showing a similar synaptic competition, consistent with the results previously reported^41^.

**Fig. 3.**
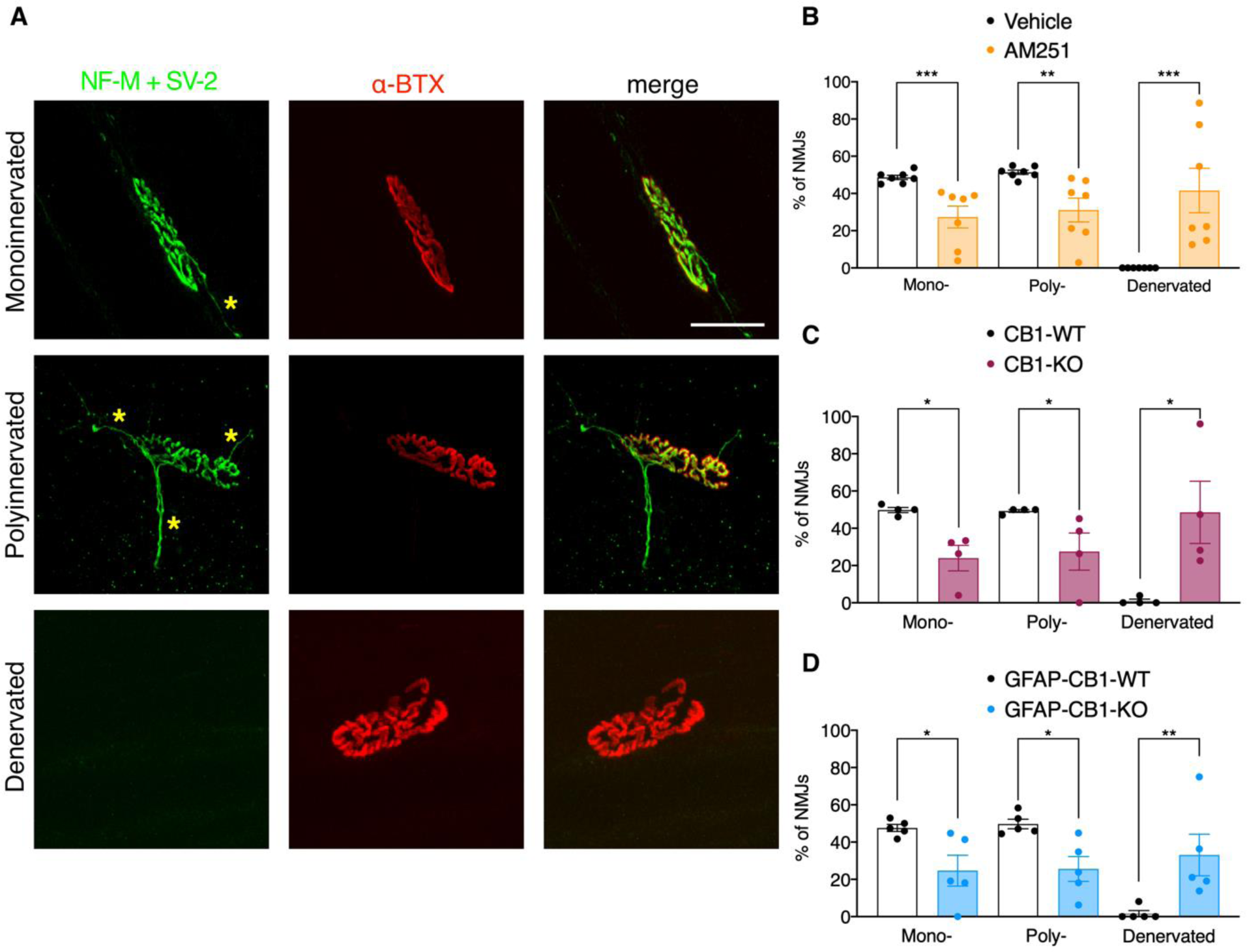
CB_1_R blockade in PSCs impairs NMJ reinnervation. (A) Representative images of the different innervation states. Number of axons has been indicated with yellow asterisks. (B) AM251 treatment decreases mono-(*** p < 0.001; N = 7, n = 65) and poly-innervated NMJs (** p < 0.01; N = 7; n = 72) with a significant increase of denervated NMJs (*** p < 0.001; N=7, n = 94). (C) Full CB1-KO mice (* p < 0.05; N = 4, n = 37) and, in particular, (D) glial-KO, GFAP-CB_1_-KO, (** p < 0.01; N = 5, n = 53) also showed enhanced denervation. Scale bar = 30 *μ*m. Results are presented as mean ± SEM.

Morphological analysis performed using YFP-Thy1 mice showed that *in vivo* treatment with AM251 (Fig. 3A, B), decreased mono-(*** p < 0.001; N = 7, n = 65) and poly-innervated NMJs (** p < 0.01; N = 7; n = 72) and increased denervated NMJs (*** p < 0.001; N=7, n = 94) compared to control, vehicle-injected mice. To confirm that the effect of the treatment was due to the block of CB_1_R we performed the same experiment in full-CB_1_-KO. Reinnervation was also greatly impaired in full CB1-KO mice with a significant increase of denervated NMJs, confirming the AM251 results (Fig. 3C, * p < 0.05; N = 4, n = 37). These results are consistent with a delayed reinnervation following nerve injury when CB_1_R were blocked.

As mentioned above, PSCs are central elements regulating NMJ maintenance, function and repair after nerve injury^34,41^. Hence, we tested if the reinnervation process was impaired when CB_1_Rs were selectively knocked out in PSCs. We found a greater denervation in GFAP-CB_1_-KO compared to GFAP-CB_1_-WT (Fig. 3D, ** p < 0.01; N = 5, n = 53), demonstrating that the effects observed with the treatment and in the full-CB_1_*-*KO were mainly mediated by CB_1_R on PSCs. These data highlight that CB_1_R on PSCs are responsible for the reinnervation processes to properly be executed.

The deleterious impact of the antagonist suggests that CB_1_R are important for the process of reinnervation to take place. Hence, this opens a new unexplored path where CB_1_R could be activated to support a better and faster reinnervation. To test this hypothesis, we used the same experimental plan described above for the antagonist and tested whether WIN 55,212-2, a CB_1_R agonist, could facilitate reinnervation.

NMJ morphological analysis indicated that the treatment with WIN 55,212-2 (0.3 mg/kg) facilitated the reinnervation as shown by a faster and more complete reinnervation with a significant increase of mono-innervated NMJs (Fig. 4A; *** p < 0.001; N=8, n = 124) and decrease of poly-innervated NMJs (Fig. 4A; *** p < 0.001; N = 8, n = 77). In another set of experiments, we determined that the 3 mg dose did not ameliorate reinnervation, with mono-(Fig. 4B; ns; N = 7, n = 124) and poly-innervation comparable to vehicle mice (ns; N = 8, n = 92). Hence, our data suggest that CB_1_R positively regulates NMJ reinnervation after 16 DPI.

**Fig. 4.**
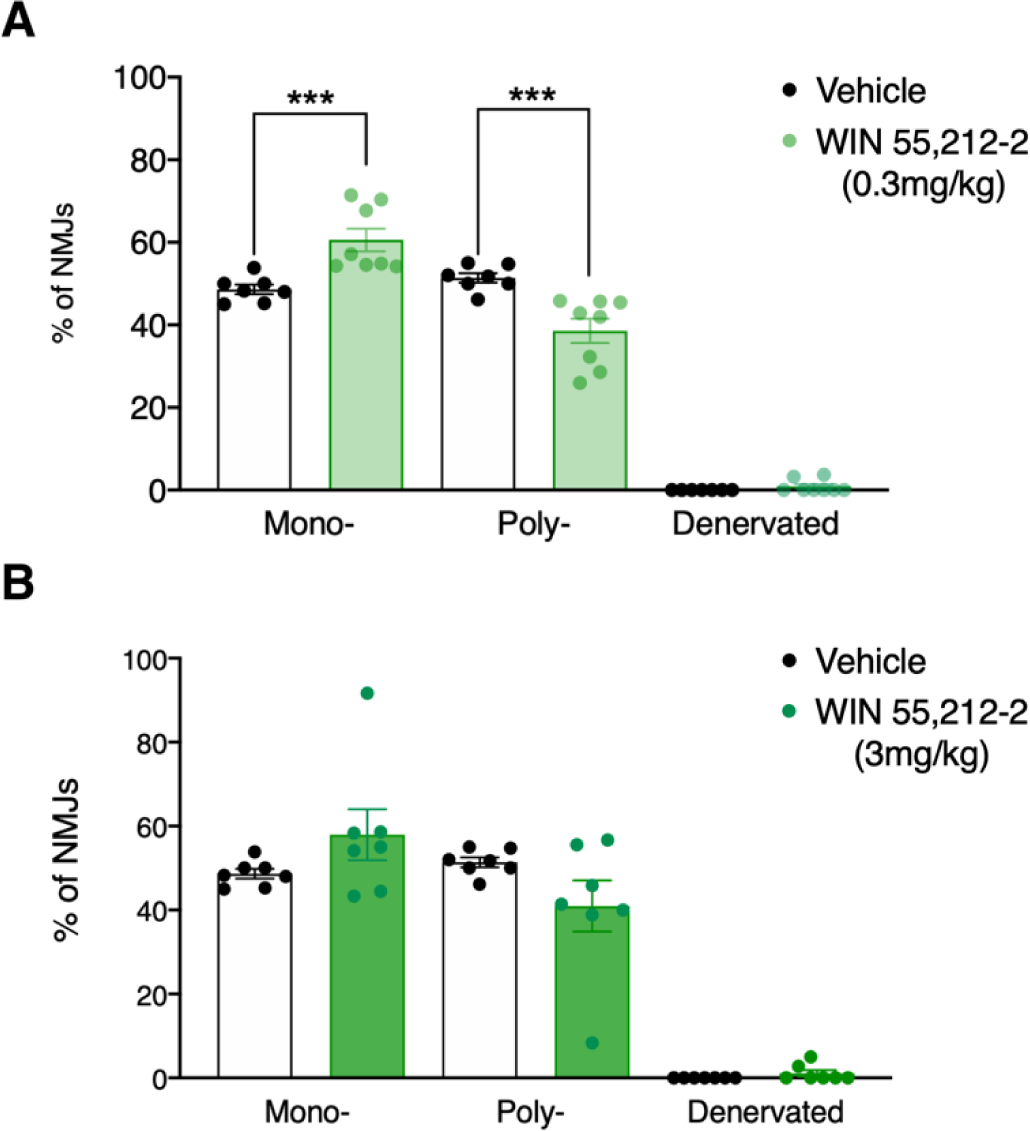
CB_1_R agonist promotes NMJ reinnervation. (A) WIN 55,212-2 (0.3 mg/kg) improved reinnervation at 16 DPI, a significant increase of mono-innervated NMJs (*** p < 0.001; N = 8, n = 124) and decrease of poly-innervated NMJs (*** p < 0.001; N = 8, n = 77). (B) In another set of experiments, 3 mg of WIN 55,212-2 dose did not alter reinnervation, with mono-(ns; N = 7, n = 124) and poly-innervation comparable to vehicle treated mice (ns; N = 8, n = 92). Results are presented as mean ± SEM.

## 4. Discussion

In this study, we established that CB_1_R is not only present at NMJs and muscle fibers^23,33^ but it is also expressed by PSCs, where it regulates denervation and reinnervation following nerve injury. We showed that the absence of the CB_1_R during the denervation stage accelerated the full NMJs denervation process that was followed by a greatly impaired reinnervation following axonal injury. Importantly, treatment with a CB_1_R agonist facilitated reinnervation.

Mechanisms regulating the reinnervation process following nerve injury are based on those involved in postnatal NMJ formation occurring during development^42^. Indeed, in both cases, NMJs are first poly-innervated and then undergo a period of synaptic competition, which ends when NMJs become mono-innervated^42,64,65^ with the elimination of the weakest nerve terminal depending on the synaptic activity^66,67^. PSCs are the main actors on the NMJ development and response to injury^34,50,68^, switching into a repair phenotype in a mAChRs-mediated process^41,50^. While different morphological and physiological changes have been observed^41,50,69^, mechanisms controlling nerve injury response through PSCs are poorly defined.

Here, we not only demonstrate that the blocking of CB_1_R (AM251-treated and in full-CB_1_-KO mice) is detrimental for nerve regeneration with an increase of denervated NMJs during the period of synaptic competition but, that the regulation is mainly, if not exclusively, mediated by PSCs as revealed using the glial-specific strategy with the GFAP-CB_1_-KO. This further suggests that PSCs response to injury was impaired when the CB_1_R was absent or altered. A delay in the NMJ reinnervation rather than a complete blockade following glial CB_1_R absence is consistent with the concept that multiple parallel mechanisms regulate NMJ reinnervation to ensure nerve regeneration.

These observations unravel a novel CB_1_R, glia-mediated, regulation of injury response at NMJs. For instance, NMJs can experience injury and repair cycles during lifetime, altering the innervation status and the muscle functions. Although peripheral nerve injuries may not be life threatening *per se*, they can be the primary outcome of different neuromuscular disease and can considerably decline the patient’s quality of life. Moreover, it has been estimated that at the slow rate of 1 mm/day regeneration rate in humans, several years will be needed for regenerating nerves to reach muscles leading to muscle atrophy and fat tissue accumulation^43^. Thus, a suitable therapeutic strategy should seek to promote neural outgrowth and accelerate motor axon regeneration. Consistent with this possibility, activation of CB_1_R by the agonist WIN55,212-2 considerably improved NMJs reinnervation. Also, study in a model of spinal nerve ligation of neuropathic pain, a WIN55,212-2 anti-hyperalgesic and anti-allodynic effects is blocked by the CB_1_R antagonist, but not the CB_2_R demonstrated that the analgesic effect is mediated by CB_1_Rs^70^. All these results indicate that the endocannabinoid system through CB_1_R may have therapeutic potential not only in the reduction of neuropathic pain but also in the improvement of motor recovery.

In addition to the novelty of a glial CB_1_R mechanism, these observations open a new avenue of possible original pharmacological opportunity to sustain, accelerate and improve motor nerve-muscle repair following injury and eventually, in neuromuscular diseases. For instance, in a disease-driven research, PSC dysfunctions prevented them to switch to repair mode caused by a gradual increased sensitivity of PSC acetylcholine muscarinic receptors (mAChRs) towards disease onset in Amyotrophic lateral sclerosis (ALS)^46^. Interestingly, the impact of CB_1_R on NMJ denervation and repair is very reminiscent of the data we obtained in ALS studies^71–73^, while CB_1_R regulates the activity of mAChRs^74,75^. The activation of the CB_1_R system of PSCs could improve their response to NMJ denervation and injury, resulting in a better balance between NMJ maintenance and repair. Moreover, several ongoing clinical trials using cannabinoid drugs are underway to test for treatment of spasticity and other muscle-debilitating symptoms^76^. The increased usage of cannabinoids in different medical conditions^77^ highlights the importance of elucidating CB_1_R functions to better understand repair of motor injury and neurodegenerative disorders.

## Acknowledgments

The authors wish to thank the different members of the laboratory who provided insightful inputs throughout the course of this project. R.R. and R.P. were supported by the Canadian Institutes of Health Research (CIHR grants #PJT-185884 and PJT-190295). R.P. was also supported by FRQS (Fonds de recherche du Québec-Santé) and held a Scientific Scholar ‘ALS Scholars in Therapeutics’ by the Sean M. Healey & AMG Center for ALS at Massachusetts General Hospital (MGH). This work was also supported by Inserm (to G.M. and L.B.); EU–FP7 (PAINCAGE, HEALTH-603191, to G.M.); the European Research Council (Micabra, ERC-2017-AdG-786467, to G.M.); Fondation pour la Recherche Medicale (DRM20101220445, to G.M., and ARF20140129235, to L.B.); French State/Agence Nationale de la Recherche (LABEX BRAIN ANR-10-LABX-43, to G.M. and JCJC MitoCB1-fat, to L.B.) and the French government in the framework of the University of Bordeaux’s IdEx “Investments for the Future - France 2030” program / GPR BRAIN_2030 (to G.M. and L.B.).

## Conflict of interest

The authors declare no conflict of interest.

## Author Contributions

Roberta Piovesana and Richard Robitaille conceived and designed the project. Luigi Bellocchio and Giovanni Marsicano provided the Full- and GFAP-CB_1_-KO, discussed the results and helped with the experimental design and data interpretation. Roberta Piovesana, Sophie Charron, Danielle Arbour acquired the data. Roberta Piovesana, Sophie Charron analyzed and interpreted the data. Roberta Piovesana and Richard Robitaille wrote the manuscript.

## Data Availability statement

The data that support the findings of this study are available from the corresponding author upon reasonable request.

## Notes

### Competing Interest Statement

The authors have declared no competing interest.

