## Supplementary material for "Cannabinoid type-1 (CB_1_) receptors in glial cells promote neuromuscular junction repair following nerve injury": Supp Table 1 and Figure 1

**Table 1. Type of innervation at 18 h post-injury.**

| Type of NMJ innervation | Definition | Example |
|---|---|---|
| **Early ongoing denervation** | Coverage of the pre- (NF-M/SV-2) on the post-synaptic endplate (α-BTX) was still present.<br><br>Less than 30% of the denervated NMJ area (orange square). | 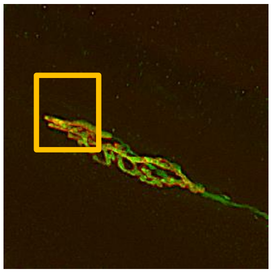 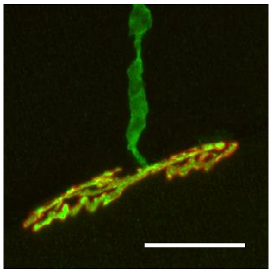 |
| **Denervating NMJ** | Up to 70% of the denervated NMJ area.<br><br>Axon is still present at the NMJs. | 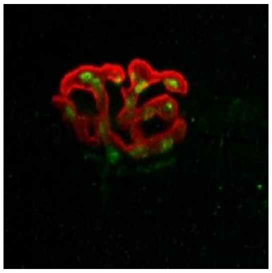 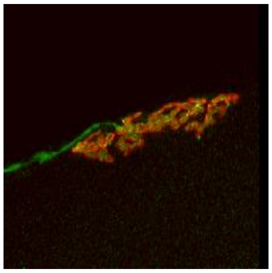 |
| **Denervated NMJ** | 70% or more of denervated NJM area.<br><br>Axon staining is absent. Some pre-synaptic labelling can still be present (yellow triangle). | 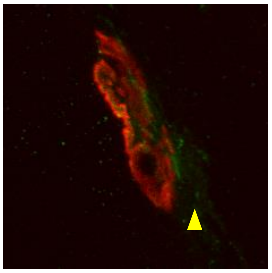 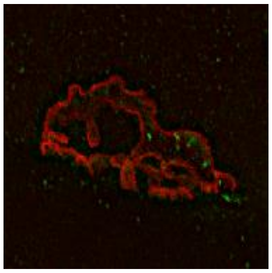 |

**Table 2. Reinnervation type at 16 days post-injury.**

| Type of Innervation | Definition | Example |
|---|---|---|
| **Mono-innervated** | Complete coverage of the pre- (NF-M/SV-2) on the post-synaptic endplate (α-BTX).<br>Coverage is made by a single axon (white arrow). | 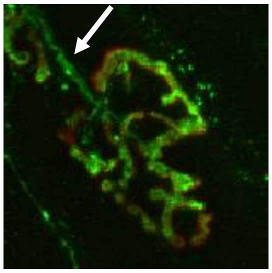 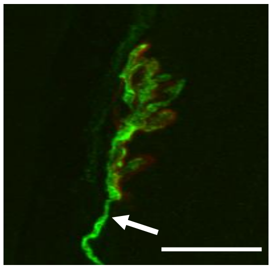 |
| **Poly-innervated** | Complete coverage of the pre- (NF-M/SV-2) and post-synaptic endplate (α-BTX).<br>Coverage is made by at least two independent axons (white arrows). | 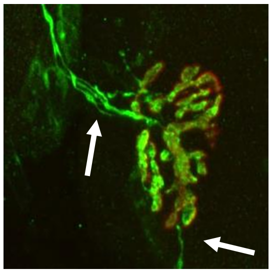 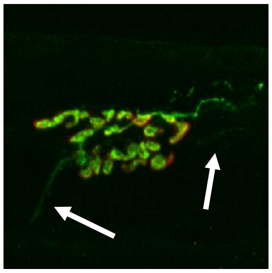 |
| **Denervated** | No nerve terminal label (NF-M/SV-2) was found at the post-synaptic endplate (α-BTX). | 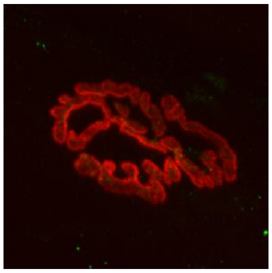 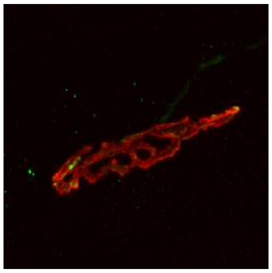 |

**Supplemental Data 1**

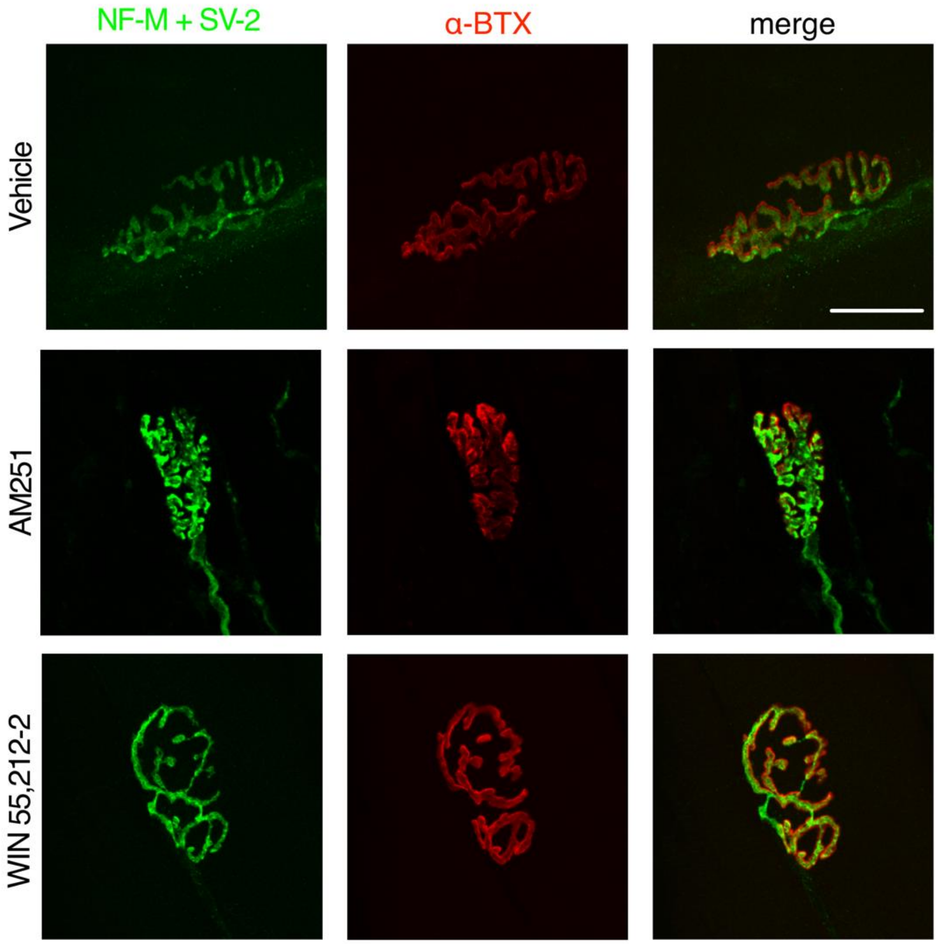
